# Cerebral Bases and Neural Dynamics of Audiovisual Temporal Binding Window: a TMS study

**DOI:** 10.1101/2025.10.01.679698

**Authors:** Solène Leblond, Tutea Atger, Isabelle Berry, Franck-Emmanuel Roux, Céline Cappe, Robin Baurès

## Abstract

The temporal binding window (TBW) refers to the time interval within which two stimuli, typically visual and auditory, are perceived as synchronous. Neural bases underlying this process consistently implicate a large-scale network with superior temporal sulcus (STS) as a central hub, alongside contributions from primary auditory and visual cortices and higher-order areas including prefrontal and posterior parietal cortex.

This study aimed to provide causal evidence for the involvement of superior temporal gyrus (STG) and intraparietal sulcus (IPS) in the simultaneity judgment (SJ) task using MRI-guided transcranial magnetic stimulation (TMS). In particular, we aimed to clarify temporal dynamics of these regions in audiovisual synchrony perception.

Forty adults performed an SJ task in which they had to determine whether stimuli were synchronous or asynchronous. Single-pulse TMS was applied over bilateral IPS, STG, or the vertex at different delays following stimulus offset.

Early stimulation of left IPS and right STG increased the proportion of “synchronous” responses. In contrast, later stimulation of bilateral STG was associated with reduced synchrony perception.

These findings provide causal evidence for a dynamic interplay between parietal and temporal regions in audiovisual temporal integration. Early IPS and STG involvement facilitates temporal integration, while later STG activity promotes perceptual segregation.

## Introduction

We live in a dynamic multisensory world where combining information from different sensory modalities is essential for a unified perception. Multisensory integration (MSI) refers to the process by which information from different senses is merged. A key principle guiding MSI is the temporal law: for two stimuli to be integrated, they must be perceived as simultaneous [1].

The Temporal Binding Window (TBW) is the temporal interval during which two visual and auditory stimuli are integrated into one event. It is typically studied using the Simultaneity Judgment (SJ) task, where participants judge whether audiovisual (AV) pairs are synchronous or asynchronous [2–5]. When both auditory first (A–V) and visual-first (V–A) stimuli are tested, a Gaussian curve is typically fitted to the percentage of “synchronous” responses across stimulus onset asynchronies (SOAs). The point of subjective simultaneity (PSS) corresponds to the SOA with the highest synchrony perception. One study reported a PSS of approximately 47 ms during an SJ task, with V–A stimulus [2]. Another found a PSS of 19 ms (V–A) when AV stimuli were spatially aligned, increasing to 32 ms when misaligned [5]. Across studies, the PSS is rarely centered at 0 ms (i.e., when auditory and visual stimuli are physically simultaneous), but often shifted toward SOAs where the visual stimulus precedes the auditory one, suggesting that asynchrony is harder to detect when the visual stimulus leads. This is consistent with auditory stimuli being processed faster, which aligns with the faster neural conduction speed compared to visual stimuli. Despite this slight bias toward visual-leading SOAs, synchrony perception remains high at 0 ms, with more than 90% of trials typically judged synchronous [2,5–9]. In the present study, participants performed an SJ task using only auditory-first pairs to facilitate task performance.

It is also possible to compute the synchrony threshold, defined as the SOA at which synchrony and asynchrony responses are equally likely (i.e., 50% “synchronous”), which is often considered the TBW’s boundaries. Researchers typically use psychometric functions rather than Gaussian fits in studies testing only one asynchrony direction (A–V or V–A).

The neural bases of MSI are now well documented. MSI relies on a distributed network spanning early sensory areas to higher-order associative and frontal regions. It can occur at low levels of processing, with evidence of direct crossmodal influences in primary sensory cortices, and extends to higher-level regions such as the prefrontal cortex, which may exert top-down control over perceptual integration [10–12]. In particular, numerous studies consistently highlighted the superior temporal sulcus (STS) as a central hub for AV integration [13,14]. The TBW, which reflects the temporal dimension of MSI, has also been the focus of extensive research. The present study aims explicitly to investigate the neural mechanisms underlying AV asynchrony detection, a key component of temporal MSI. An fMRI study identified a predominantly left-lateralized network, including the left anterior superior temporal gyrus, left inferior parietal cortex, left medial frontal gyrus, and right operculum [15]. In contrast, a literature review reported no consistent lateralization and described a broad network that includes the auditory and visual cortices, the dorsal fronto-parietal attention system, and key MSI regions such as the superior colliculi, insula, inferior parietal cortex, and again the STS [9,16,17].

However, these findings rely on correlational methods (EEG, fMRI). Causal evidence comes from a study using transcranial direct current stimulation (tDCS), which showed that anodal stimulation of the right posterior parietal cortex (rPPC) during an SJ task significantly narrowed the TBW, reducing “synchronous” responses at 150 ms SOA from 77.1% to 53.96% [18]. Such a result demonstrates that modulating rPPC activity can influence synchrony perception.

Although EEG is a correlational method, it provides valuable insight into the temporal dynamics of neural activation. High-density EEG has revealed distinct activation patterns for A–V and V–A trials across three key time windows: early (39–95 ms), intermediate (142–222 ms), and late (297–351 ms) [6]. Gamma-band phase synchronization between 170 and 250 ms is associated with asynchronous perception, whereas low-frequency synchronization (23–28 Hz) occurs earlier (40–70 ms) during synchronous trials [19]. Later brain activity linked to asynchrony detection was confirmed by increased ERP positivity between 210 and 270 ms [7]. These findings suggest distinct temporal signatures: early neural responses linked to synchrony detection, while later activity reflects asynchrony processing.

To expand on these findings, the present study used single**-**pulse TMS, a causal method, to investigate the role of the intraparietal sulcus (IPS) and superior temporal gyrus (STG) during an SJ task. Anatomical MRI allowed guiding the stimulation with high spatial precision. Since TMS transiently disrupts neural activity [20,21], performance changes would indicate the targeted region’s causal involvement. Previous studies have shown that TMS over these regions can modulate performance in cognitive tasks [22–26].

Given that TMS effects last less than 50 ms [20], applying pulses at different delays after stimulus onset may selectively interfere with synchrony or asynchrony detection. TMS thus provides a unique tool to probe both the neural structures and temporal dynamics supporting AV simultaneity perception.

The present study aims to confirm the involvement of the IPS and STG in the TBW using an SJ task and to identify the timing of their contribution by applying single**-**pulse TMS at different post**-** stimulus delays.

## Materials and methods

### Participants

Forty-one healthy participants were recruited (mean age ± SD: 23.75 ± 2.28 years; range: 21-29 years; 15 females). One participant discontinued the task due to discomfort from the stimulation, resulting in a final sample of 40 participants with normal or corrected-to-normal vision. All participants were naïve to the experimental procedure and the purpose of the study. The study was approved by the French Ethical “Comité de Protection des Personnes” (IRB number: 2023-A01271-44). All individuals provided written informed consent, in accordance with ethical guidelines.

## MRI-Guided Neuronavigation and TMS Parameters

### Inclusion visit and anatomical MRI

Each participant underwent a medical visit to confirm eligibility for MRI and TMS. A high-resolution T1-weighted anatomical scan was then acquired (3T Philips; flip angle = 8°; TR = 8.1 ms; TE = 3.7 ms; FOV = 240 × 240 mm; voxel size = 1 mm^3^) to guide stimulation.

### Neuronavigation system

Frameless stereotactic neuronavigation (Brainsight, Rogue Research, Montreal, Canada) was used. Reflective markers attached to the participant’s head and the TMS coil allowed real-time tracking. Co-registration with individual MRIs was achieved via digitized landmarks (nasion, tragus).

Target regions, left/right intraparietal sulcus (IPS), superior temporal gyrus (STG), and vertex (VTX), were identified precisely and marked on individual cortical reconstructions by neurosurgeon Prof. Roux. The IPS was defined as 1 cm posterior to the junction of the postcentral sulcus along the sulcus, and the STG was defined as the posterior portion of the superior temporal gyrus, approximately 1 cm behind the postcentral sulcus (Figure 1).

**Figure 1.**
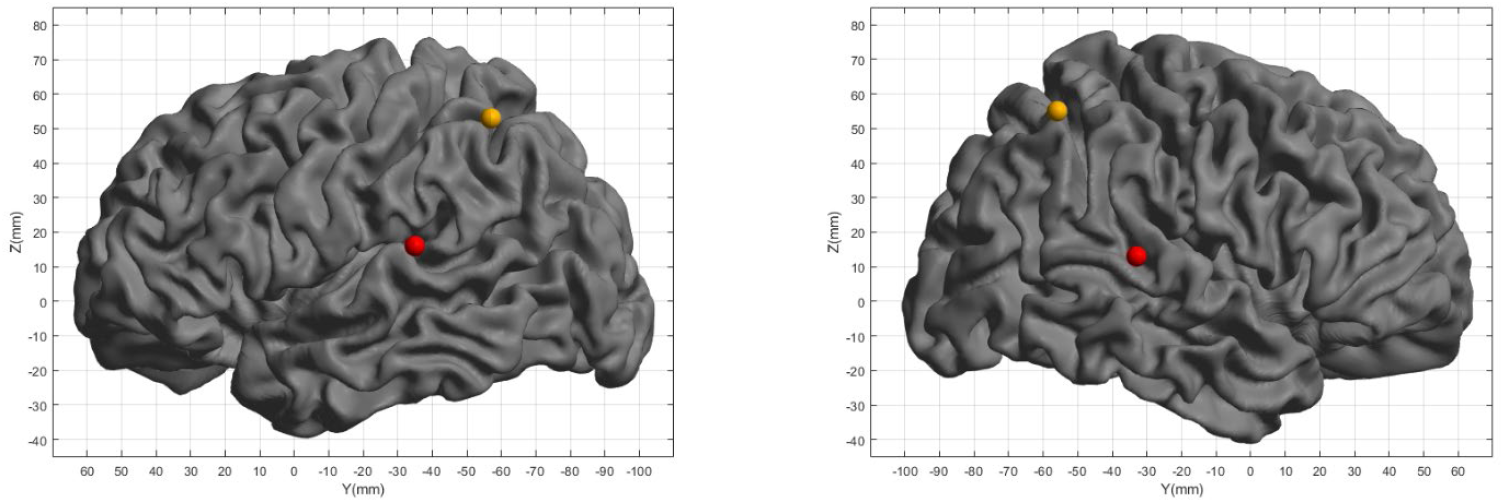
TMS stimulation sites in left and right hemispheres. Mean stimulation sites in the intraparietal sulcus (IPS, orange points) and superior temporal gyrus (STG, red points) are shown for the left (left panel) and right (right panel) hemispheres. The IPS was defined as 1 cm posterior to the junction of the postcentral sulcus along the sulcus. The STG was defined as the posterior portion of the superior temporal gyrus, approximately 1 cm behind the postcentral sulcus. MNI coordinates (x, y, z ± SD) across participants were: Right IPS, 33 ± 4, –56 ± 4, 55 ± 4 mm; Left IPS, –36 ± 3, –57 ± 5, 53 ± 7 mm; Right STG, 61 ± 3, –33 ± 7, 13 ± 6 mm; Left STG, –55 ± 20, –35 ± 5, 16 ± 25 mm.

Stimulation lateralization was randomly assigned across participants, forming two distinct groups. The mean location of the TMS site at the right IPS in MNI space was at 33 ± 4; –56 ± 4; 55 ± 4 mm (x; y; z ± SD). The mean location of the TMS site at the left IPS was –36 ± 3; –57 ± 5; 53 ± 7 mm. The mean location of the right STG was 61 ± 3; –33 ± 7; 13 ± 6 mm, and the left STG was – 55 ± 20; –35 ± 5; 16 ± 25 mm (Figure 1). The vertex location was 0.4 ± 2; –30 ± 6; 83 ± 9 mm. See Supplementary Material for individual participants’ coordinates.

The VTX served as a control site. While some control conditions use subthreshold stimulation or a reversed coil to mimic the auditory click without cortical activation [25], these can lead to perceptible differences in sound intensity or tactile sensation, compromising participant blinding, especially in tasks involving auditory stimuli. Therefore, it is preferable to use a control brain area that can be either a brain region known to be uninvolved in the task [23,27] or a neutral anatomical location such as the VTX. Indeed, previous TMS/fMRI studies have shown that stimulation over the vertex has minimal influence on ongoing brain processes [28,29]. Given the distributed nature of the multisensory integration network, the VTX was chosen as a neutral site to reduce the risk of inadvertently stimulating task-relevant areas.

### Transcranial Magnetic Stimulation (TMS)

Single-pulse TMS was delivered using a Magstim Rapid2 with a 70 mm figure-of-eight coil (focal field ~1.5–3 cm^2^) [30,31]. Stimulation intensity was set at 100% of each participant’s resting motor threshold (RMT), determined as the minimal output eliciting a visible hand twitch in ≥50% of 10 trials [32,33]. Mean intensity was 61% ± 6% (range 47–70%) for IPS and vertex; for comfort, STG intensity was reduced to 75% of this value (mean 46% ± 5%, range 35–57%). The coil was held tangentially with a posterior-to-anterior orientation, optimizing stimulation efficiency [30].

### Simultaneity Judgment task

#### Simultaneity Judgment (SJ) Task protocol

Participants performed an SJ task indicating whether an auditory and a visual stimuli were perfectly synchronous or delayed (Figure 2). The auditory stimulus was a 50 ms, 2000 Hz pure tone, delivered via Etymotic ER4-XR wired headphones. The visual stimulus was a 50 ms white circle displayed on a 25.6-inch monitor (1440 × 1080 pixels; 50 × 40 cm), connected to a Dell Precision 5820 Tower desktop (Intel Xeon 3.80 GHz). A variable SOA was introduced between the two stimuli.

**Figure 2.**
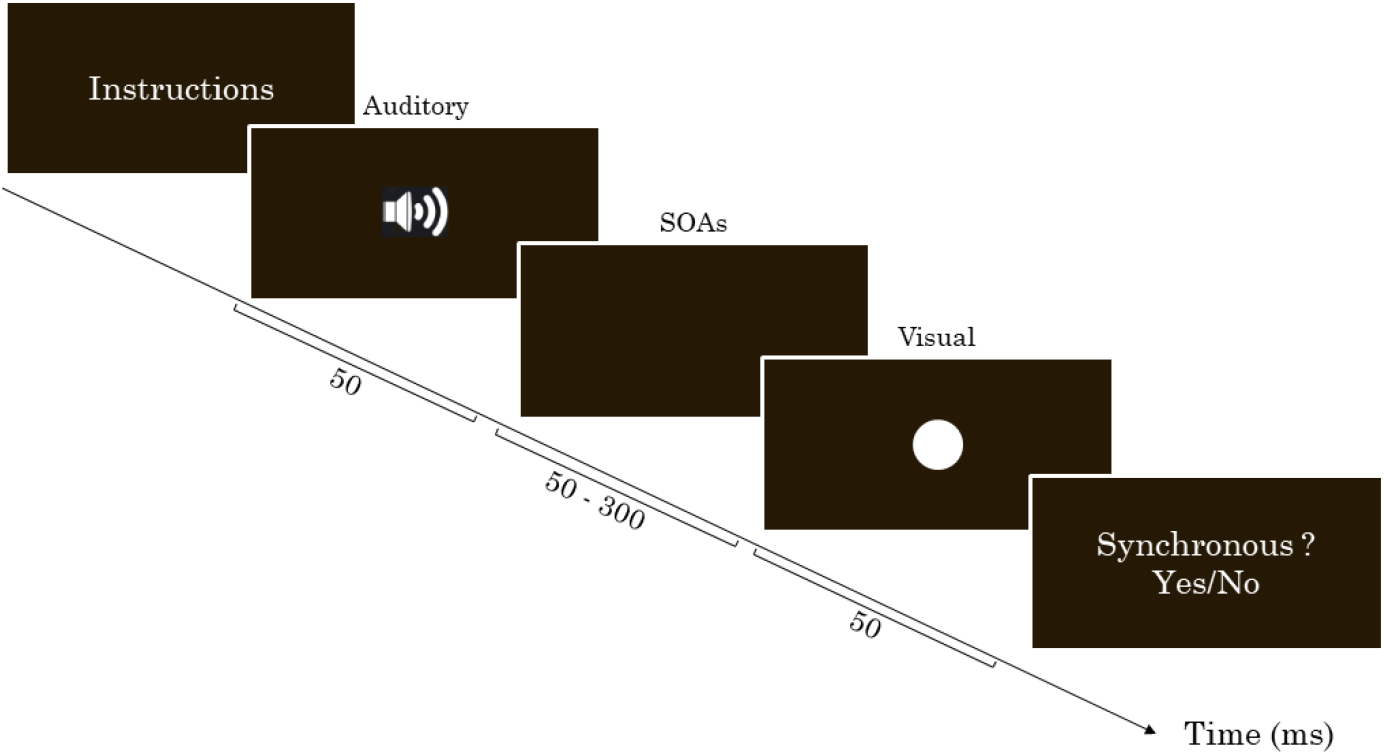
Trial structure of the Simultaneity Judgment (SJ) Task. Illustration of audiovisual stimulus presentation and participant response.

#### Pre-test phase

Each participant completed a pre-test session without stimulation. Participants performed an SJ task using 12 SOAs ranging from 50 to 400 ms, presented across six blocks of 36 trials (12 SOAs × 3 repetitions), for a total of 216 randomized trials. For each trial, they reported via keyboard whether the stimuli were synchronous or asynchronous, using the hand ipsilateral to the future stimulation site to avoid TMS-induced motor interference. The TMS coil was placed away from the scalp to expose participants to the stimulation noise without actual brain stimulation.

Pre-test data were used to derive individual psychometric curves showing the percentage of “synchronous” responses across SOAs (Figure 3). From this, five personalized delays (SOAs corresponding to 60%, 70%, 75%, 80%, and 90% synchrony) were selected for the main TMS session.

**Figure 3.**
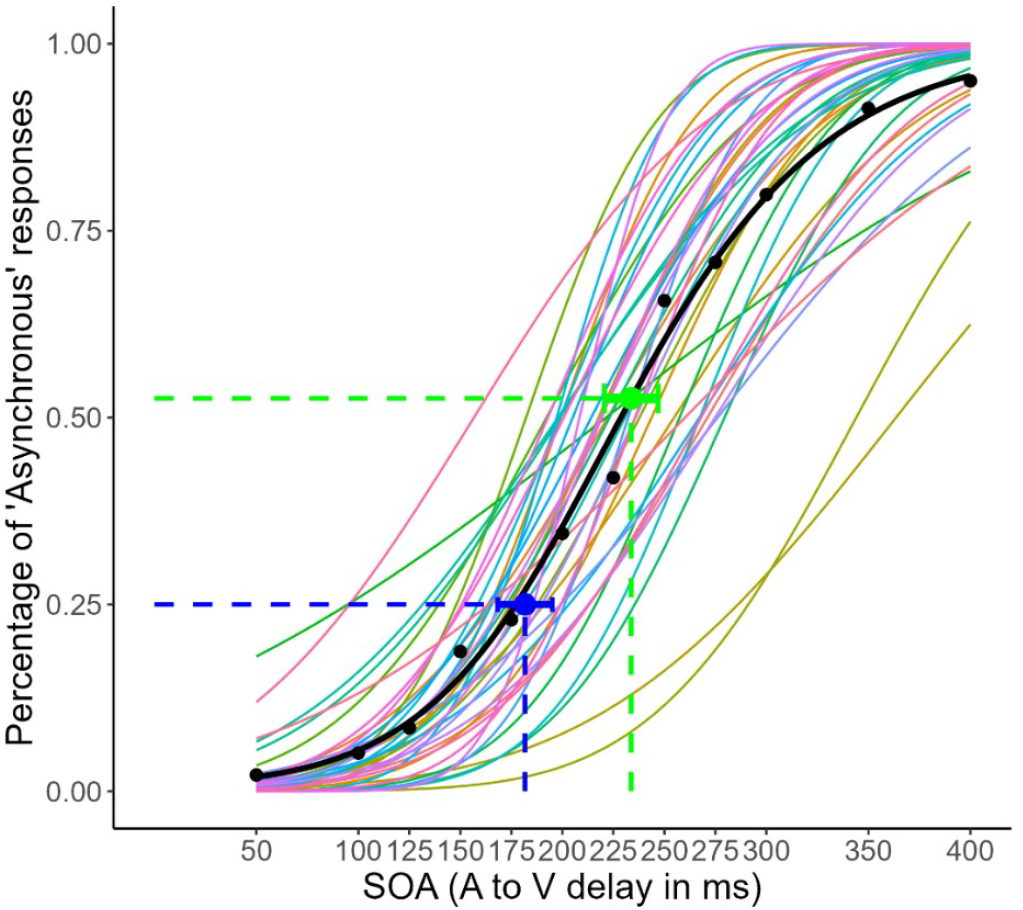
Percentage of asynchronous response as a function of SOA for the 38 participants. Each curve represents one participant. Note that we used in the TMS session SOAs corresponding to 60, 70, 75, 80, and 90% of synchronous response, therefore in this figure corresponding to the SOA leading to 40, 30, 25, 20 and 10% of asynchronous response. Black dots represent the mean percentage of asynchronous responses averaged across participants for each SOA. The black curve represents the mean psychometric function fitting this averaged dataset. The simultaneity threshold and its 95% confidence interval are shown in green, and the SOA corresponding to 25% asynchronous responses and its CI_95_ are displayed in blue.

#### Tests with TMS stimulation

The TMS session followed the pre-test and used each participant’s individualized SOA corresponding to 75% synchrony as the main condition. Single TMS pulses were delivered at this SOA after one of six post-stimulus delays (50, 100, 150, 200, 250, 300 ms) to probe the temporal dynamics of the targeted region. Non-stimulated filler trials (SOAs at 60%, 70%, 80%, and 90% synchrony) were included to prevent anticipation and maintain variability.

Each block included 30 trials: 18 with stimulation (3 per pulse delay) and 12 non-stimulated filler trials (4 SOAs × 3). Participants completed 12 blocks per stimulation site, for a total of 360 trials per site, including 216 with TMS. Three stimulation targets were tested: IPS, STG, and VTX. The stimulated hemisphere (left or right) was randomly assigned, and site order was counterbalanced across participants.

Before and after the TMS session, participants completed six standardized tests assessing visuospatial attention, motor coordination (upper/lower limbs), balance, fine motor skills, and auditory abilities, to verify the absence of TMS-induced side effects. Additionally, participants completed the Edinburgh Handedness Inventory [34] to assess their right-handedness (mean laterality quotient = 81.9 ± 20.7).

### Data Analysis

#### Pre-Test: Psychometric curve

Psychometric curves were fitted for each participant based on the percentage of “synchronous” responses across AV SOAs. From these fits, we extracted the SOAs corresponding to 60%, 70%, 75%, 80%, and 90% “synchronous” responses, with only the 75% SOA used for TMS stimulation trials. The simultaneity threshold (50% synchrony) was taken as a proxy for the TBW boundary. The just noticeable difference (JND) was defined as half the distance between the 25% and 75% synchrony points; smaller JNDs indicate sharper temporal sensitivity. All metrics were estimated using the R quickpsy package [35], averaged across participants, and visualized through a group-level psychometric curve (Figure 3).

A repeated-measures ANOVA was first performed to assess the effect of SOA on synchrony perception. An independent-samples t-test (assuming unequal variance) compared SOA75 between left and right hemisphere groups to control for group differences. As the number of males and females was heavily unbalanced, HRM ANOVA [36–38] was performed.

#### Test with TMS stimulation: Data Filtering

A total of 43,200 trials were conducted (360 trials × 3 stimulation sites × 40 participants). Only stimulation trials corresponding to the SOA_75_ were retained, resulting in 25,920 trials.

A first filtering step removed 2,400 trials (9.26%) in which participants responded before the TMS pulse, mainly at the longest post-stimulus delays (22.2% discarded at 300 ms), reducing the dataset to 23,520 trials. This filtering process ensured that all retained responses occurred after both the AV stimuli and the TMS pulse.

Two participants were excluded from the analysis: one for excessive early responses (over 60% discarded at 250 and 300 ms), and one due to TMS intolerance, leading to abnormally low synchrony responses across all delays (below 23% vs. the expected 75%). The final dataset included 22,467 valid trials (91.24% of usable trials) from 38 participants, equally split between left and right hemisphere stimulation groups (n = 19 per group).

#### Test with TMS stimulation: Statistical Analysis

An initial repeated-measures ANOVA was performed on the Percentage of “synchronous” responses, with Zone and TMS Pulse Delay as within-variables and Hemisphere as a between variable. Pairwise t-tests were used, with Bonferroni correction applied at each TMS pulse delay level to account for multiple comparisons.

In a subsequent analysis, we tested whether the difference in the percentage of “synchronous” responses between STG vs. Vertex and IPS vs. Vertex significantly differed from zero at each TMS pulse delay separately. We performed Student’s t-tests with Bonferroni correction applied at each TMS pulse delay level to account for multiple comparisons. Additionally, 95% confidence intervals (CI_95_) were computed for all differences. All reported *p*-values in the Results section are Bonferroni-adjusted.

## Results

To investigate the neural bases of the temporal binding window (TBW), participants first completed a pre-test session of the simultaneity judgment (SJ) task, followed by a session in which MRI-guided TMS was applied over the STG and IPS at different post-stimulus intervals.

### Pre-test session

During the pre-tests, participants performed an SJ task. For each participant, the psychometric curve representing the percentage of asynchronous responses as a function of SOA was generated (Figure 3).

The mean simultaneity threshold (SOA_50_) across the 38 participants was 234 ms, CI_95_ = [220: 247]. The JND had a mean value of 77 ms, with a 95% confidence interval [68: 86]. The mean SOA_75_, corresponding to 75% of “synchronous” responses, was 182 ms, with a 95% confidence interval of [168: 195]. A repeated-measures ANOVA confirmed a significant effect of SOA on the percentage of “synchronous” responses, F(11, 407) = 347.56, 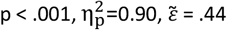.

The control tests showed that the two Hemisphere groups did not differ in the SOA_75_, t(25.8) = 1.38, p =.181, nor did Sex influence the SOA_75_, F(1, 35.46) = 1.24, p =.273, as shown by the HRM ANOVA. However, the absence of proof is no proof of absence, so we performed equivalence tests using the TOSTER package [39,40]. Assuming that the SOA_75_ could present an equivalence boundary of at most 25% without implying a significant difference, the equivalence tests for both Sex and Hemisphere supported the acceptance of the null hypothesis.

### Tests with TMS stimulation

Our initial ANOVA investigated possible differences in the “synchronous” response rate among the three stimulation sites (IPS, GTS, and Vertex) and two hemispheres (right and left), at the different TMS pulse delays. The ANOVA showed a significant effect of TMS Pulse Delay, which however turned non-significant after sphericity violation correction, *F*(5, 180) = 2.36, *p* =.08, 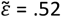. TMS pulse delay and Zone interacted, *F*(10, 360) = 3.92, *p* <.001, 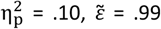. However, probably due to a substantial correction for multiple comparisons, no pairwise difference was found in the post-tests. No other significant effect was found.

In a second analysis, we compared the difference in percentage of “synchronous” responses during IPS or STG stimulation and the control site (VTX) stimulation, at each TMS pulse delay separately, using paired t-tests (Figure 4).

**Figure 4.**
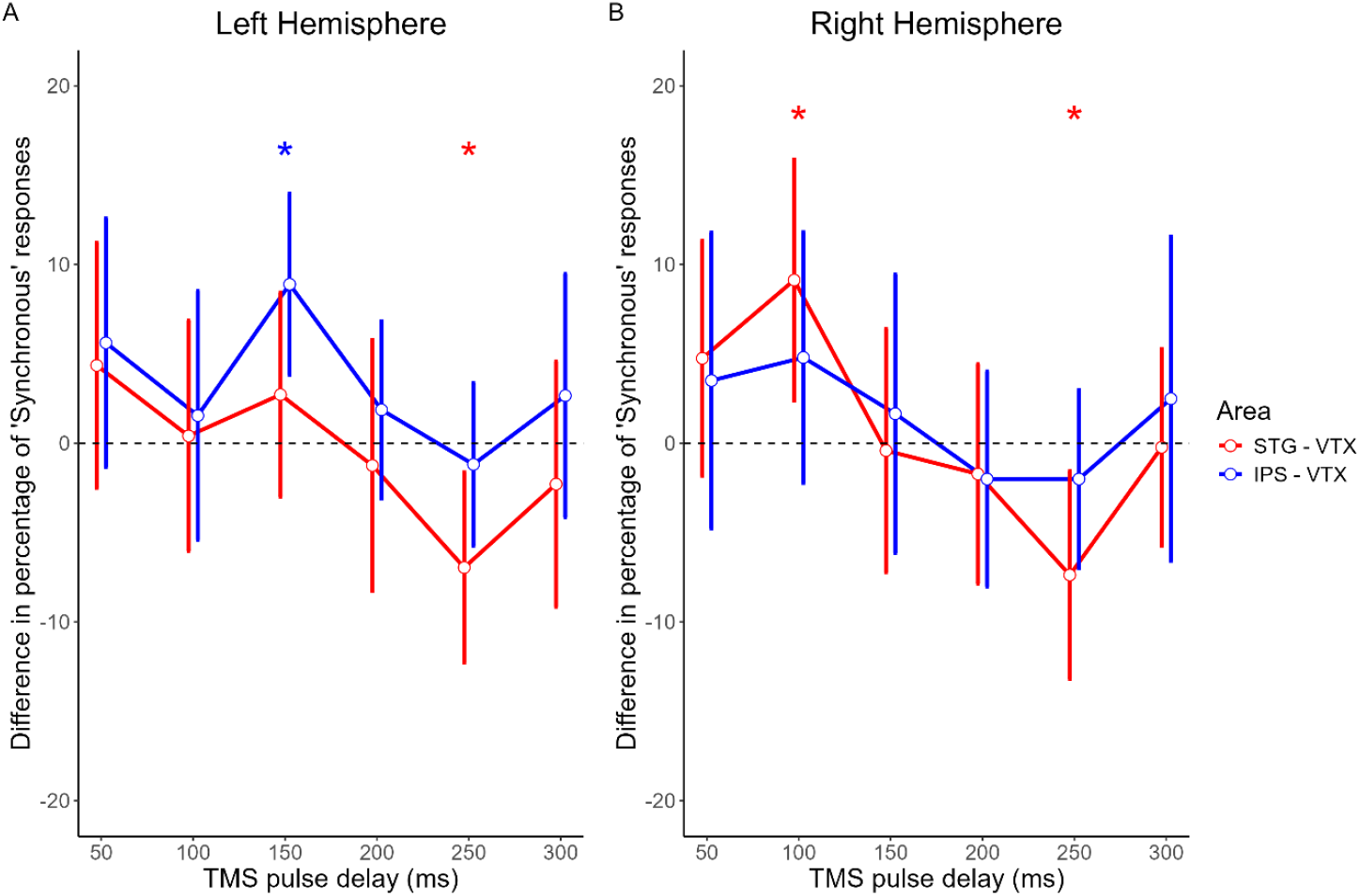
Mean differences in the percentage of “Synchronous” responses between stimulation sites (IPS = Intraparietal Sulcus, STG = Superior Temporal Gyrus) and the control site (Vertex = VTX), as a function of TMS pulse delay after stimulus offset, for the left (A) and right (B) hemispheres. Error bars represent the 95% confidence interval of the difference. Note that if the error bars do not cross the zero line, the difference is significantly different from zero (at an uncorrected p-value threshold of.05).

Significant effects were observed at specific stimulation sites and TMS delays. In the left hemisphere, the difference in “synchronous” responses between IPS and VTX stimulation at the 150 ms TMS pulse delay was positive, indicating that IPS stimulation led to a significantly higher percentage of “synchronous” responses compared to stimulation of the VTX (t(18) = 3.68, *p* =.003, mean difference of percentage = 8.90%, CI_95_ = [3.82: 14.00]). The effect size was high (Cohen’s d_z_ = 0.84), indicating that this stimulation effect was consistently observed across most participants. In contrast, the difference between STG and VTX stimulation at a 250 ms TMS pulse delay was negative, meaning that STG stimulation led to a significantly lower percentage of “synchronous” responses relative to the vertex condition (t(18) = 2.75, p =.027, mean difference of percentage = −6.96%, CI_95_ = [−12.30: −1.63]). The effect size was medium (d_z_ = 0.63), suggesting a moderately consistent effect across participants.

In the right hemisphere, the difference between STG and VTX stimulation at 100 ms TMS pulse delay was positive, indicating a significantly higher percentage of “synchronous” responses during STG stimulation compared to VTX stimulation (t(18) = 2.84, p =.022, mean difference of percentage = 9.13%, CI_95_ = [2.38: 15.90]). The effect size was medium (d_z_ = 0.65), indicating a moderately consistent increase across participants. Conversely, at 250 ms, the difference between STG and VTX was negative meaning that STG stimulation induced a significant lower percentage of “synchronous” responses compared to VTX stimulation (t(18) = 2.67, p =.031, mean difference of percentage = −7.38%, CI_95_ = [−13.20, –1.57]) with a medium effect size as well (d_z_ = 0.61). At any delay, no significant differences in the percentage of “synchronous” responses were found between IPS and VTX stimulation.

## Discussion

The present study investigated the causal involvement of the STG and the IPS in AV synchrony perception, and by extension, their roles in the TBW. To this end, participants performed an SJ task while TMS was applied to these regions in both hemispheres separately. Our results revealed a significant contribution of the left IPS and bilateral STG to AV synchrony perception. Furthermore, by delivering single-pulse TMS at different pulse delays following stimulus offset, the study provided new insights into the temporal dynamics of these regions. Specifically, we showed early engagement of the left IPS and right STG, which was associated with increased synchronicity perception, and later, the contribution of the bilateral STG was associated with decreased synchrony perception.

Participants completed a pre-test session without stimulation to determine their individual SOA_75_ for the TMS session. This also allowed us to estimate the average simultaneity threshold reflecting the point at which participants are equally likely to perceive AV stimuli as synchronous or asynchronous, which was 234 ms, higher than typically reported in the literature. For example, Zampini et al. reported a mean threshold of 161 ms when the stimuli were presented at the same spatial location [5]. Their threshold of 161 ms was obtained under conditions where auditory and visual stimuli were spatially coincident. This raises the question of whether, in our study, participants perceived the stimuli as spatially aligned, even though the auditory signal was delivered through headphones. Kaganovich and Schumaker [7] distinguished “good” and “poor” performers, exhibiting a mean simultaneity threshold of 134 ms and a substantially wider threshold of 270 ms, respectively. In the present study, we did not classify participants according to their performance levels, which could partly account for differences in the simultaneity thresholds observed. Stevenson and Wallace [41] reported a 173 ms mean threshold, significantly varying with stimulus nature (speech, flash-beep, and tool sounds). This suggests threshold variability across studies may also stem from differences in stimulus characteristics such as duration, intensity, or complexity. Overall, the simultaneity threshold observed in our study is consistent with the existing literature, which highlights that various factors influence this measure. Differences in analysis methods, stimulus properties, experimental parameters, and equipment can all contribute to variability in threshold values, emphasizing the multifactorial nature of AV temporal perception.

Participants then completed the TMS session. A repeated-measures ANOVA on “synchronous” responses revealed a significant main effect of TMS Pulse Delay, but it did not remain significant after correction for sphericity violations. An interaction between TMS Pulse Delay and Zone was also observed, but post hoc comparisons failed to reveal significant effects, likely due to stringent multiple comparison correction. The apparent lack of effect on the raw data suggests that the effects of TMS over the different sites are too subtle to withstand statistical test corrections, and a more fine-tuned analysis appears necessary. To limit the number of comparisons, which may obscure potential differences, we conducted a second analysis comparing the difference in the percentage of “synchronous” responses between IPS and control site stimulation (IPS–VTX), and between STG and VTX stimulation (STG–VTX), using paired t-tests with Bonferroni correction applied at each Pulse Delay separately.

First, these tests revealed a significant difference between IPS and VTX stimulation in the left hemisphere at the 150 ms delay, but no involvement of right IPS. This aligns with fMRI findings, reporting left-lateralized activation in the left inferior parietal cortex [15] but also contradicts Zmigrod and Zmigrod [18], who used TDCs and found right-hemisphere lateralization, with rPPC stimulation narrowing the TBW by approximately 30%. The discrepancies may arise from methodological differences: tDCS applies continuous low-intensity electrical stimulation while TMS produces brief magnetic pulses to induce neural activity. In their study, tDCS was applied continuously for 15 minutes, whereas we used single TMS pulses at various post-stimulus delays. Therefore, the absence of TMS effects at specific delays may reflect a temporal mismatch, where stimulation missed the critical time window during the AV temporal binding process. Thus, right IPS involvement cannot be excluded but might have been missed due to suboptimal stimulation timing. Future studies should use shorter pulse intervals or using rTMS to enhance the chances to stimulate during the appropriate period.

The present study also revealed a significant difference between STG and VTX stimulation in the right hemisphere at the 100 ms delay, and bilateral effects at 250 ms. While Lux et al. [15] reported left-lateralized activation, their findings specifically highlighted the left anterior STG, potentially overlapping with our STG stimulation site. The STS, often described as the MSI hub [17], is anatomically and functionally connected to the STG. A beep-flash task study showed increased BOLD responses in both STS and STG under synchronous conditions, with bilateral activation [42]. In a subsequent study using speech stimuli [43], they observed comparable STS involvement with spatially distinct subdivisions within the STS, with a central “synchrony region” flanked by “asynchrony regions”. These findings support the STS as a central, bilaterally active hub for AV (a)synchrony perception across stimulus types.

A similar pattern emerged for both targeted brain areas: TMS applied to IPS and STG led to more “synchronous” responses than VTX stimulation at early delays (around 100 ms), followed by fewer at around 250 ms. This pattern suggests early stimulation may have enhanced synchrony perception, effectively broadening the TBW, whereas later stimulation increased sensitivity to asynchrony, narrowing the TBW. These results align with previous findings, showing early neural activity (from 40 ms) is associated with AV synchrony detection, whereas later responses (200 ms) relate to asynchrony processing [7,19]. This temporal dissociation highlights distinct neural mechanisms for synchrony vs. asynchrony perception.

Previous research has also highlighted the role of frontal regions, notably the dorsal fronto-parietal attention network, in synchrony perception [44]. In a review, Zhou et al. proposed a network model in which the STS serves as a core hub for temporal AV processing, with the concomitant implication of frontal areas including the insula and prefrontal cortex [45,46]. Although the present study focused on STG and IPS, future work may aim to probe frontal contributions to AV asynchrony detection via TMS to elucidate the broader network dynamics further.

## Acknowledgments

This research was supported by the Tremplin call for proposals from the University of Toulouse. We are grateful to the Brainsight technical team for their assistance with neuronavigation and to Laurie Galas for her guidance and training in using neuronavigation and TMS. We also thank Pr. Isabelle Berry, the study investigator, for her support during participant inclusion visits, and Nathalie Vayssière for her help in preparing the CPP submission. We acknowledge Maxime Rosito for his technical and IT assistance. Finally, we thank the Inserm/UPS UMR1214 Technical Platform and the IRM Tonic platform for their assistance with MRI acquisitions.

## Availability of data and materials

Data that supports the findings of this study is available on this link: https://osf.io/ach2n/?view_only=e8673372d24a4e5891c2d5d38d6ee233

## Author contributions

- **Conceptualization:** Solène Leblond, Céline Cappe, Robin Baurès
- **Data curation:** Solène Leblond, Tutea Atger, Robin Baurès
- **Formal analysis:** Solène Leblond, Robin Baurès
- **Investigation:** Solène Leblond, Tutea Atger, Isabelle Berry
- **Methodology:** Solène Leblond, Franck-Emmanuel Roux, Céline Cappe, Robin Baurès
- **Project administration:** Solène Leblond, Isabelle Berry, Céline Cappe, Robin Baurès
- **Resources:** –
- **Software:** Solène Leblond (experiment code), Solène Leblond & Robin Baurès (R data analysis scripts)
- **Supervision:** Franck-Emmanuel Roux, Céline Cappe, Robin Baurès
- **Validation:** Solène Leblond, Céline Cappe, Robin Baurès
- **Visualization:** Solène Leblond, Céline Cappe, Robin Baurès
- **Writing – original draft:** Solène Leblond
- **Writing – review & editing:** Solène Leblond, Franck-Emmanuel Roux, Céline Cappe, Robin Baurès

## Competiting interests

The authors declare no competiting interests.

## Declaration of generative AI and AI-assisted technologies in the writing process

Generative AI and AI-assisted technologies were used to improve the readability and language of the manuscript. All content generated by the model was reviewed and edited by the authors for accuracy and clarity.

