## Supplementary Tables for "Cerebral Bases and Neural Dynamics of Audiovisual Temporal Binding Window: a TMS study"

### Supplementary Material

| N° | Participant | Hemisphere | X (mm) | Y (mm) | Z (mm) | Area |
|---|---|---|---|---|---|---|
| 1 | UN-001 | Right | 66,3 | -25,8 | 7,67 | STG |
| 2 | EE-005 | Right | 64,06 | -30,09 | 11,34 | STG |
| 3 | BL-011 | Left | -64,52 | -37,2 | 12,04 | STG |
| 4 | OU-014 | Right | 60,44 | -22,13 | 9,79 | STG |
| 5 | EA-018 | Left | -64,71 | -35,37 | 16,7 | STG |
| 6 | EO-010 | Right | 56,83 | -34,88 | 10,37 | STG |
| 7 | AU-016 | Left | -64,58 | -25,75 | 15,38 | STG |
| 8 | AY-012 | Left | -63,49 | -38,18 | 7,54 | STG |
| 11 | AA-007 | Left | -57,87 | -35,14 | 14,01 | STG |
| 12 | EE-006 | Right | 61,63 | -37,36 | 12,24 | STG |
| 13 | UG-009 | Left | -58,1 | -32,58 | 0,82 | STG |
| 14 | AE-015 | Right | 57,82 | -36,47 | 14,7 | STG |
| 15 | OA-020 | Right | 62,41 | -22,17 | 3,71 | STG |
| 16 | AA-008 | Right | 66,48 | -36,04 | 16,19 | STG |
| 17 | AE-013 | Left | -0,8 | -29,63 | 92,54 | STG |
| 18 | EE-035 | Right | 60,74 | -43,54 | 11,71 | STG |
| 19 | LN-024 | Left | -62,36 | -40,96 | 14,23 | STG |
| 20 | AO-042 | Left | -60,78 | -24,17 | 4,27 | STG |
| 22 | OE-004 | Right | 59,26 | -28,62 | 8,41 | STG |
| 23 | HO-037 | Left | 1,36 | -37,3 | 82,98 | STG |
| 24 | LV-039 | Left | -61,98 | -27,83 | 4,53 | STG |
| 25 | GU-040 | Left | -59,6 | -41,54 | 3,9 | STG |
| 26 | UI-029 | Right | 57,8 | -39,56 | 21,46 | STG |
| 27 | OA-023 | Left | -63,42 | -37,5 | 11,79 | STG |
| 28 | UL-022 | Left | -63,71 | -44,07 | 12,64 | STG |
| 29 | OA-034 | Right | 61,57 | -33,29 | 8,52 | STG |
| 30 | AU-027 | Left | -62,82 | -34,98 | 15,58 | STG |
| 31 | AU-036 | Right | 60,45 | -31,98 | 3,28 | STG |
| 32 | HM-032 | Left | -54,1 | -34,35 | 14,8 | STG |
| 33 | UA-030 | Right | 55,4 | -37,71 | 19,13 | STG |
| 34 | AX-031 | Left | -58,68 | -28,99 | 13,35 | STG |
| 35 | AI-021 | Right | 60,86 | -28,34 | 5,61 | STG |
| 36 | AA-019 | Right | 60 | -18,75 | -3,6 | STG |
| 37 | II-041 | Left | -63,43 | -34,48 | 11,83 | STG |
| 38 | OO-038 | Right | 60,27 | -42,15 | 10,24 | STG |
| 39 | HU-026 | Right | 59,77 | -44,86 | 16,04 | STG |
| 40 | AE-028 | Left | -66,23 | -35,45 | 18,56 | STG |
| 41 | EA-025 | Right | 60,03 | -38,82 | 11,77 | STG |

Table 1: Individual coordinates of the superior temporal gyrus (STG) stimulation sites for all participants (19 left hemisphere, 19 right hemisphere). Coordinates are reported in MNI space (x, y, z) for each participant.

| N° | Participant | Hemisphere | X (mm) | Y (mm) | Z (mm) | Area |
|---|---|---|---|---|---|---|
| 1 | UN-001 | Right | 34,4 | -56,94 | 52,99 | IPS |
| 2 | EE-005 | Right | 39,43 | -51,38 | 55,27 | IPS |
| 3 | BL-011 | Left | -32,23 | -53,27 | 60,28 | IPS |
| 4 | OU-014 | Right | 32,03 | -59,69 | 56,25 | IPS |
| 5 | EA-018 | Left | -38,63 | -57,35 | 62,65 | IPS |
| 6 | EO-010 | Right | 36,18 | -58,19 | 47,54 | IPS |
| 7 | AU-016 | Left | -30,16 | -49,4 | 68,77 | IPS |
| 8 | AY-012 | Left | -33,17 | -61,99 | 59,2 | IPS |
| 11 | AA-007 | Left | -37,44 | -52,35 | 55,22 | IPS |
| 12 | EE-006 | Right | 36,87 | -58,1 | 51,68 | IPS |
| 13 | UG-009 | Left | -35,51 | -67,47 | 42,07 | IPS |
| 14 | AE-015 | Right | 33,09 | -52,32 | 54,17 | IPS |
| 15 | OA-020 | Right | 40,68 | -46,89 | 58,19 | IPS |
| 16 | AA-008 | Right | 34,91 | -61,12 | 57,17 | IPS |
| 17 | AE-013 | Left | -40,45 | -52,56 | 53,17 | IPS |
| 18 | EE-035 | Right | 29,57 | -56,16 | 65,79 | IPS |
| 19 | LN-024 | Left | -38,13 | -55,87 | 60,34 | IPS |
| 20 | AO-042 | Left | -29,61 | -60,16 | 50,67 | IPS |
| 22 | OE-004 | Right | 31,81 | -53,73 | 56,5 | IPS |
| 23 | HO-037 | Left | -39,49 | -55,58 | 57,11 | IPS |
| 24 | LV-039 | Left | -37,71 | -61,2 | 47,25 | IPS |
| 25 | GU-040 | Left | -32,32 | -61,53 | 53,02 | IPS |
| 26 | UI-029 | Right | 30,17 | -49,39 | 56,32 | IPS |
| 27 | OA-023 | Left | -32,41 | -61,63 | 60,73 | IPS |
| 28 | UL-022 | Left | -35,59 | -60,59 | 58,67 | IPS |
| 29 | OA-034 | Right | 31,19 | -58,9 | 52,43 | IPS |
| 30 | AU-027 | Left | -39,09 | -60,37 | 53,56 | IPS |
| 31 | AU-036 | Right | 25,35 | -57,14 | 60,12 | IPS |
| 32 | HM-032 | Left | -36,17 | -54,77 | 66,42 | IPS |
| 33 | UA-030 | Right | 30,13 | -50,43 | 58,2 | IPS |
| 34 | AX-031 | Left | -36,29 | -62,86 | 51,58 | IPS |
| 36 | AA-019 | Right | 35,96 | -60,56 | 48,86 | IPS |
| 37 | II-041 | Left | -34,99 | -52,67 | 62,14 | IPS |
| 38 | OO-038 | Right | 27,83 | -58,3 | 63,48 | IPS |
| 39 | HU-026 | Right | 32,07 | -61,19 | 54,5 | IPS |
| 40 | AE-028 | Left | -36,46 | -58,13 | 67,19 | IPS |
| 41 | EA-025 | Right | 36,21 | -56,44 | 54,9 | IPS |

Table 2: Individual coordinates of the intraparietal sulcus (IPS) stimulation sites for all participants (19 left hemisphere, 19 right hemisphere). Coordinates are reported in MNI space (x, y, z) for each participant.

| N° | Participant | Hemisphere | X (mm) | Y (mm) | Z (mm) | Area |
|---|---|---|---|---|---|---|
| 1 | UN-001 | Right | -1,13 | -31,17 | 92,32 | Vertex |
| 2 | EE-005 | Right | 0,5 | -32,31 | 86,81 | Vertex |
| 3 | BL-011 | Left | 4,12 | -26,75 | 96,31 | Vertex |
| 4 | OU-014 | Right | -0,63 | -31,63 | 98,78 | Vertex |
| 5 | EA-018 | Left | 1,48 | -28,34 | 105,48 | Vertex |
| 6 | EO-010 | Right | -1,7 | -31,59 | 95,61 | Vertex |
| 7 | AU-016 | Left | -0,85 | -37,73 | 101,67 | Vertex |
| 8 | AY-012 | Left | -2,5 | -23,32 | 98,78 | Vertex |
| 11 | AA-007 | Left | 1,04 | -33,89 | 94,35 | Vertex |
| 12 | EE-006 | Right | 0,72 | -31,74 | 91,24 | Vertex |
| 13 | UG-009 | Left | 4,58 | -39,65 | 91,57 | Vertex |
| 14 | AE-015 | Right | -0,35 | -28,09 | 96,27 | Vertex |
| 15 | OA-020 | Right | -1,81 | -32,73 | 102,85 | Vertex |
| 16 | AA-008 | Right | -0,69 | -35,12 | 97,37 | Vertex |
| 17 | AE-013 | Left | -0,8 | -29,63 | 92,54 | Vertex |
| 18 | EE-035 | Right | 0,63 | -32,39 | 87,49 | Vertex |
| 19 | LN-024 | Left | 3,57 | -29,19 | 85,45 | Vertex |
| 20 | AO-042 | Left | -0,64 | -35,14 | 69,87 | Vertex |
| 22 | OE-004 | Right | 2,13 | -28,26 | 92,4 | Vertex |
| 23 | HO-037 | Left | 1,36 | -37,3 | 82,89 | Vertex |
| 24 | LV-039 | Left | 0,14 | -18,77 | 79,18 | Vertex |
| 25 | GU-040 | Left | 1,69 | -29,57 | 80,46 | Vertex |
| 26 | UI-029 | Right | 4,05 | -23,88 | 78,03 | Vertex |
| 27 | OA-023 | Left | -0,77 | -28,96 | 86,77 | Vertex |
| 28 | UL-022 | Left | -1,73 | -30,15 | 84,48 | Vertex |
| 29 | OA-034 | Right | 0,7 | -24,91 | 78,34 | Vertex |
| 30 | AU-027 | Left | 1,41 | -27,91 | 85,96 | Vertex |
| 31 | AU-036 | Right | -0,74 | -35,36 | 76,54 | Vertex |
| 32 | HM-032 | Left | 0,79 | -30,1 | 89,59 | Vertex |
| 33 | UA-030 | Right | 1,67 | -18,98 | 80,06 | Vertex |
| 34 | AX-031 | Left | 2,9 | -32,11 | 81,56 | Vertex |
| 35 | AI-021 | Right | -2,2 | -11,37 | 81,03 | Vertex |
| 36 | AA-019 | Right | -3,1 | -31,41 | 91,89 | Vertex |
| 37 | II-041 | Left | 0,92 | -30,58 | 72,61 | Vertex |
| 38 | OO-038 | Right | -0,75 | -27,61 | 75,12 | Vertex |
| 39 | HU-026 | Right | 1,73 | -24,19 | 86,35 | Vertex |
| 40 | AE-028 | Left | 0,47 | -39,83 | 99,85 | Vertex |
| 41 | EA-025 | Right | 1,06 | -32,64 | 79,65 | Vertex |

Table 3: Individual coordinates of the Vertex (VTX) stimulation sites for all participants (19 left hemisphere, 19 right hemisphere). Coordinates are reported in MNI space (x, y, z) for each participant.
